# Erythroid spectrin binding modulates peroxidase and catalase activity of heme proteins

**DOI:** 10.1101/2021.09.06.459089

**Authors:** Dipayan Bose, Shantanu Aggarwal, Debashree Das, Chandrabhas Narayana, Abhijit Chakrabarti

## Abstract

Hemoglobin oxidation due to oxidative stress and disease conditions leads to generation of ROS (reactive oxygen species) and membrane attachment of hemoglobin *in-vivo*, where its redox activity leads to peroxidative damage of membrane lipids and proteins. Spectrin, the major component of the RBC membrane skeleton, is known to interact with hemoglobin and, here this interaction is shown to increase hemoglobin peroxidase activity in the presence of reducing substrate ABTS (2’, 2’-Azino-Bis-3-Ethylbenzothiazoline-6-Sulfonic Acid). It is also shown that in the absence of reducing substrate, spectrin forms covalently cross-linked aggregates with hemoglobin which display no peroxidase activity. This may have implications in the clearance of ROS and limiting peroxidative damage. Spectrin is found to modulate the peroxidase activity of different hemoglobin variants like A, E, and S, and of isolated globin chains from each of these variants. This may be of importance in disease states like sickle cell disease and HbE-β-thalassemia, where increased oxidative damage and free globin subunits are present due to the defects inherent in the hemoglobin variants associated with these diseases. This hypothesis is corroborated by lipid peroxidation experiments. The modulatory role of spectrin is shown to extend to other heme proteins, namely catalase and cytochrome-c. Experiments with free heme and Raman spectroscopy of heme proteins in the presence of spectrin show that structural alterations occur in the heme moiety of the heme proteins on spectrin binding, which may be the structural basis of increased enzyme activity.

## Introduction

Spectrin is the major component of the membrane skeleton in mature human erythrocytes (RBCs) [1]. It has been shown that spectrin interacts with hemoglobin, as evidenced by fluorescence and light scattering based studies, as well as proteomic investigations [2–5]. So far, the only known consequence of the spectrin-hemoglobin interaction is via the ability of spectrin to act as a chaperone for hemoglobin [2, 6, 7].

Spectrin has been shown to act as a chaperone for hemoglobin, and display a preference for more structurally unstable isoforms of hemoglobin, like hemoglobin E (HbE), and such hemoglobin interactions are known to be redox active, as seen in the oxidative cross-linking of HbE and spectrin [7–9]. Spectrin is also able to act as a chaperone for other heme containing proteins like HRP, and in that case it is seen that there is a modulation of enzyme activity of HRP as well [10]. A recent proteomic study from our lab has identified several redox regulators and heme containing proteins like cytochrome, catalase, peroxiredoxin etc as a major part of the interactome of spectrin [3]. Thus, from all these lines of evidence we believe spectrin may have a role in the redox biology of hemoglobin and other heme containing proteins [7].

Moreover, it is known that spectrin can accept liberated heme, that is lost from hemoglobin due to stress conditions, and this spectrin-heme conjugate may then show modulated the redox activity [11].

Thus, the known association of spectrin with redox active heme containing proteins, as well as its ability to bind hemoglobin motivates us to investigate whether spectrin association has an effect on the redox activity of hemoglobin. This becomes especially important considering the important biological role of hemoglobin and its well documented involvement in redox biology.

It is well known that auto-oxidation of hemoglobin takes place under *in-vivo* conditions to generate met-hemoglobin [Fe(III)], peroxide, and superoxide radicals, collectively known as reactive oxygen species (ROS) [12–14]. Under normal conditions a certain basal level of hemoglobin oxidation and corresponding level of ROS is always present in RBCs [15]. However, under certain conditions such as, under drug induced stress, hypoxic stress, oxidative stress or under disease states the rate of hemoglobin auto-oxidation and ROS generation is greatly increased [16].

Abnormal hemoglobin variants are known to have increased oxidative susceptibility and increased ROS production, such as in the case of HbE, which is known to be unstable and often associated with β-thalassemia [17, 18]. Thalassemic conditions are known promote the formation of free globin monomers, which then generate hemichromes and cause membrane oxidation and increased ROS and oxidative stress [19–21]. Another hemoglobin variant, HbS, associated with sickle cell disease is known to cause conditions where a fraction of oxidised hemoglobin remains attached to the RBC membrane where it causes lipid oxidative damage and membrane abnormalities [22, 23].

Moreover, membrane bound hemoglobin is found in complexes with membrane-skeletal proteins such as band 4.2, ankyrin and spectrin [24], and similar type of spectrin-hemoglobin association, as evidenced by decrease in thiol groups of spectrin, is noted in model β-thalassemia systems [20]. Thus, it is seen that the pathology of hemoglobin diseases is manifested by the membrane attachment of hemeoglobins and free hemes and the resultant lipid peroxidative damage and protein oxidation; for example, as seen in sickle cell disease and thalassemia [20, 25].

Thus, given the importance of the redox activity of membrane attached hemoglobin, its known association with spectrin [24], and the known ability of spectrin to take part in the redox biology of heme proteins [2, 5, 8–10], we have investigated the role of spectrin binding in the peroxidase and catalase activity of heme proteins and free heme.

In the present work we have determined the effect of spectrin interaction on the peroxidase activity of HbA, HbE and HbS as well as their isolated subunits. We have also checked the modulation of catalase activity on spectrin interaction for RBC resident heme protein catalase. Also, we have studied modulation of peroxidase activity of non-RBC resident cytochrome-c. Moreover, to gain added conformational insight we have performed Raman spectroscopy for spectrin bound heme proteins and heme.

## Experimental procedures

ABTS, catalase from horse muscle, cytochrome-c from bovine heart, MgCl_2_, PMSF, EDTA, p-hydroxymercuribenzoic acid, DTT, thiobarbituric acid (TBA), trichloro-acetic acid (TCA), sodium hydrogen phosphate, (heme) hemin chloride, NaOH, acetic acid, Tris, Sephadex G-100, Sepharose CL-4B, DEAE cellulose and CM cellulose were purchased from Sigma. All water used for experiments was purified via a Millipore system.

Stock solutions of cytochrome-c and catalase were made in 100mM phosphate buffer, pH 6.8 and 50mM Tris-HCl, pH 8.0 respectively. heme was prepared in 0.01N NaOH. Concentrations for cytochrome-c, catalase and heme were determined spectrophotometrically using molar absorbances of 106000 M^−1^ cm^−1^ (409nm), 324000 M^−1^ cm^−1^ (405nm) and 58000 M^−1^ cm^−1^ (385 nm) respectively [26–28].

### Isolation and purification of dimeric erythroid spectrin

Dimeric erythroid spectrin was isolated from clean white RBC ghost membranes following protocol elaborated in earlier studies from our lab [29–31]. Briefly, ovine blood was collected from animals slaughtered for commercial purposes, RBCs were pelleted by centrifugation, washed with PBS, lysed in hypotonic lysis buffer and RBC membranes were collected by centrifugation and contaminating hemoglobin was removed by washing with lysis buffer to yield clean white ghosts. Spectrin was removed from ghost membranes and further purified by 30% ammonium sulphate precipitation and run through Sepharose CL-4B column to give final pure product. Purity of preparation was checked by 8% SDS PAGE analysis and concentration of preparation was determined from known O.D. of 10.7 at 280 nm for 1% spectrin solution [32]. Spectrin preparation was stored at −20°C for a maximum of one month.

### Isolation and purification of HbA, HbE and HbS

Human hemoglobin variant HbA was isolated from blood samples collected from healthy volunteers with proper informed consent. Blood samples from homozygous HbE and HbS patients were obtained from Ramkrishna Mission Seva Pratisthan Hospital, Kolkata, India, with informed written consent of the patients following the guidelines of the Institutional Ethical Committee. Samples were characterized using Bio-Rad Variant HPLC system and variant hemoglobin content was determined [18].

RBCs were collected by centrifugation, washed with PBS and lysed with three volumes of 1mM Tris-HCl pH 8.0 at 4°C and lysate was purified by gel filtration on Sephadex G-100 column (45×1cm) using 5mM Tris-HCl 50mM KCl pH 8.0 buffer. Purity was checked by 12% SDS PAGE analysis and concentration was checked using molar extinction coefficient of 125000 at 415 nm. Presence of oxy form of hemoglobin was confirmed using absorbance at 415 nm and 541 nm [4]. Hemoglobin preparation was stored at −70° C for a maximum of one week.

### Isolation and purification of α and β globin subunits from hemoglobin variants

The α and β globin subunits were isolated from each of the hemoglobin variants using previously published protocol [6, 33, 34]. Briefly, 100mg PMB per 1g of hemoglobin was dissolved in minimum volume of 0.1M KOH and 1M acetic acid was added till very light precipitate persisted. PMB solution was added to 50mg/ml solution of hemoglobin in 10mM Tris-HCl 200mM NaCl, pH 8.0, and final pH was adjusted to 6.0 by addition of 1M acetic acid and mixture was incubated at 4° C for 12 hours.

PMB bound globin subunits were separated using two column selective ion exchange chromatography. α-PMB was isolated by equilibrating globin-PMB mixture with 10mM phosphate buffer pH 8.0 and passing through DEAE-cellulose column equilibrated with the same buffer. Similarly, β-PMB was isolated by equilibrating globin-PMB mixture and passing through CM-cellulose column equilibrated with the same buffer. The PMB was removed from isolated globin chains by addition of 50 mM β-mercaptoethanol in 0.1M phosphate buffer pH 7.5. Globin chains were then purified by gel filtration on BioGel P2 column. The purity was checked by 12% SDS page analysis and concentration was measured by Bradford method using BSA as standard. Globin chains were stored at 4° C for not more than 48 hours.

### Fluorescence binding study of spectrin with heme proteins

The interaction of spectrin with different heme proteins, namely, hemoglobin variants A, E and S, α and β globin subunits from each of the hemoglobin variants, catalase, and cytochrome-c were quantified in terms of the apparent dissociation constant (K_d_).

Spectrin was covalently labelled with FITC in sodium carbonate-bicarbonate buffer of pH 9.0. 2.0 mg spectrin dimer was incubated for an hour at 4° C with 50-fold molar excess of FITC, taken in a small volume of N, N-dimethylformamide. The FITC-labelled spectrin (F-spectrin) was separated and purified from the reaction mixture by gel filtration on a Sephadex G-50 column using 10mM Tris-HCl, 20mM KCl, pH 8.0. The concentration of F-spectrin was determined using Bradford method and that of fluorescein was determined spectrophotometrically from the absorbance at 495 nm using molar extinction coefficient of 76,000 M^−1^ cm^−1^. The labelling ratio of fluorescein to spectrin in F-spectrin was determined to be around 2 to 3 fluorescein per spectrin dimer [6, 35].

Steady state fluorescence measurements were performed using a Cary Varian Eclipse fluorescence spectrometer using 1cm path length quartz cuvettes with 5-nm slits for both excitation and emission channels. F-spectrin in the range of 20-50 nM was excited at 495 nm and emission was monitored from 510-600 nm. Sequential additions of different heme proteins from concentrated stock solutions were done and resultant decrease in fluorescence intensity of F-spectrin was monitored at 520 nm.

The extent of fluorescence quenching of F-spectrin as a function of increasing concentrations of different heme protein samples were analysed using the following equations [30] to give the apparent dissociation constant (K_d_) values of spectrin-heme protein interactions.

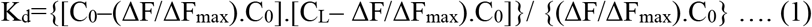

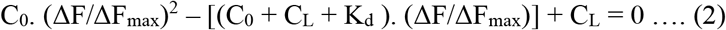

In equations (1) and (2), ΔF is the change in fluorescence emission intensity at 520 nm for each point on the titration curve and ΔF_max_ denotes the same when a given heme protein is completely bound to F-spectrin, C_L_ is the concentration of the ligand (given heme protein) at any given point in the titration curve, and C_0_ is the initial concentration of F-spectrin.

The y-axis intercept of the plot of 1/ΔF vs. 1/C_L_ gives the numerical value of 1/ ΔF_max_. In turn the calculated ΔF_max_ values can be put into equation 1 and the K_d_ values for each point on the titration curve can be generated, from which the mean K_d_ value is determined.

All experimental points for binding isotherm were fitted by least-square analysis using the Microcal Origin software package (Version 8.0) from Microcal Software Inc., Northampton, MA. The, K_d_ values are represented as mean ± standard deviation (SD) of at least 5 independent experiments.

### Peroxidase and catalase activity measurement of heme proteins

The peroxidase activity of hemoglobin variants, cytochrome-c and globin subunits from each of the hemoglobin variants were determined using the ABTS assay [10, 36, 37].

The linear range of the peroxidase activity for all the proteins and heme were determined from a plot of the initial velocity (v_0_) of the enzyme activity against the concentration of the enzyme. In case of the hemoglobin variants, isolated globin chains and cytochrome-c the enzyme activities were measured in the range of 5-30 nM, in case of catalase it was measured from 0.5-2 nM and in case of heme it was measured at 50-200 nM.

Peroxidase activity was measured in 50mM sodium-phosphate-citrate buffer pH 5.0 using 20 mM ABTS and 10 mM H2O2 in case of hemoglobin variants, isolated globin chains and cytochrome-c; in case of heme 4 mM H_2_O_2_ was used. These concentrations were chosen such that under experimental conditions reproducible measurements could be made with minimum noise. The evolution of coloured product was followed at 405 nm with time using a Cary Varian 50 Bio spectrophotometer and the initial linear region was considered for v_0_ calculations. The initial velocity of the enzyme reaction was expressed as the rate of disappearance of peroxide with time considering a stoichiometry of 2:1 ABTS coloured product formed to peroxide substrate consumed [37, 38].

Similarly, the linear range of enzyme activity of catalase was determined from a plot of v_0_ versus protein concentration in the range of 0.5-2 nM catalase (68). Enzyme activity was assayed in 50mM Tris-HCl pH 8.0 buffer using 4 mM H_2_O_2_. The rate of disappearance of peroxide with time was followed at 240nm with respect to a standard curve of H_2_O_2_ absorbance (240nm) vs. concentration determined using the permanganate titration method [39]. Initial velocity rates were expressed as rate of peroxide consumption with time.

The Michaelis-Menten constants V_max_ and K_m_ were also determined for each individual heme protein and heme.

### Effect of spectrin on enzyme activity of heme proteins

The effect of spectrin on the enzyme activities of the different heme proteins and catalytic activity of heme was determined by measuring the v_0_, K_m_ and V_max_ values of the enzyme reactions in presence of increasing concentrations of spectrin.

To determine the specificity of spectrin induced enzyme activity modulation, the activities of the different heme proteins and heme were also measured in the presence of BSA as a negative control.

The native peroxidase and catalase activity of spectrin was checked to determine its contribution to measured enzyme activity.

To differentiate whether the differences in enzyme parameters, came from the protection of heme proteins and heme from oxidative damage, or from a modulation of their enzymatic activity, the heme proteins and heme, were incubated with a given amount of spectrin in the presence of 4 mM H_2_O_2_ in the absence of ABTS and the residual enzyme activity was measured with respect to time by addition of ABTS. The initial rate of ABTS oxidation before incubation with peroxide is the initial activity of an enzyme, which is considered 100%, the initial rate of ABTS oxidation after incubation with peroxide for a given time period, in the absence of ABTS is the residual activity, which is expressed as percentage of initial activity.

Moreover, cross-linking of heme proteins to spectrin was checked under these experimental conditions according to previously published protocol [9]. Different heme proteins, 0.5 μM −1μM each were incubated with 1μM spectrin in presence of 4 mM peroxide at 37°C for 15 minutes and the resultant mixture was run on a 4% SDS gel and stained with Coomassie blue and as previously established, the band b was used for determining relative cross-linking amounts [9].

### Assay of lipid peroxidation by TBA method

Phospholipid vesicles of 100% DOPE composition were prepared using previously elaborated protocol [40]. Different combinations of 10 mM total phospholipid concentration of these prepared SUVs were taken and incubated with 10 mM hydrogen peroxide in the presence and absence of 10 nM heme proteins and 40 nM spectrin in combination with 20 mM ABTS. The extent of lipid peroxidation was measured using the TBA method according to previously published protocol using the TBA assay [41]. Briefly, 0.1 ml of sample containing 0.2 μM total lipid was incubated with 2 ml of the TBA reagent (15% w/v TCA, 0.375 w/v TBA, 0.25 N HCl) in boiling water for 15 minutes and cooled to room temperature. Precipitate was dissolved by addition of 0.2 ml of 1 N NaOH. Lipid peroxidation product malondialdehyde was assayed against a blank at 535 nm. The malondialdehyde concentration was calculated using extinction coefficient of 156000 M^−1^ cm^−1^.

### Raman spectra of heme proteins and heme in presence and absence of spectrin

Raman spectra were acquired in a Horiba LabRAM HR instrument using a 633nm laser (He-Ne laser, model 25-LHR-151-230, Melles-Griot, U.S.A.) as the excitation source. An epi-illuminated microscope (Nikon 50i, Nikon, Japan) was used both to excite the sample and collect Raman-scattered light in a backscattering geometry. Typically, a 60x infinity corrected objective (Nikon Plan Apo, Japan, NA 0.9) was used and 60 second accumulation time was used for a minimum of 2 accumulations. Raman spectra of 300 μM each of the hemoglobin variants and cytochrome-c were acquired in presence and absence of 3μM spectrin in 1mM Tris-HCl buffer pH 8.0. Similarly, the Raman spectra of heme in powder form, and in solution in presence of spectrin was acquired.

## Results

### Determination of equillibrium dissociation constants of spectrin-heme-protein interactions using fluorescein-conjugated spectrin

The heme proteins were found to interact with spectrin with apparent dissociation constants in the low micromolar range. Catalase exhibited auto-fluorescence in the experimentally monitored range and thus dissociation constants in its case could not be determined. It was seen that in general spectrin showed higher affinity for those hemoglobin variants that are known to be more structurally unstable and thus have more perturbed conformations than normal HbA [5]. This fact was further strengthened by the observation that spectrin had about the same affinity for the isolated α-globin chains from all the hemoglobin variants, which are known to have identical sequence. On the other hand, the β-globin subunits, which carry the actual mutations, from the variant hemoglobins, showed higher affinity for spectrin binding than those from HbA. Spectrin was also found to interact with cytochrome-c with a much higher affinity as compared to hemoglobin interaction. Representative fluorescence quenching curve, double reciprocal plot and binding isotherm of spectrin interaction with cytochrome-c is given in **Figure 1**. All binding data is summarized in **Table 1.**

**Figure 1:**
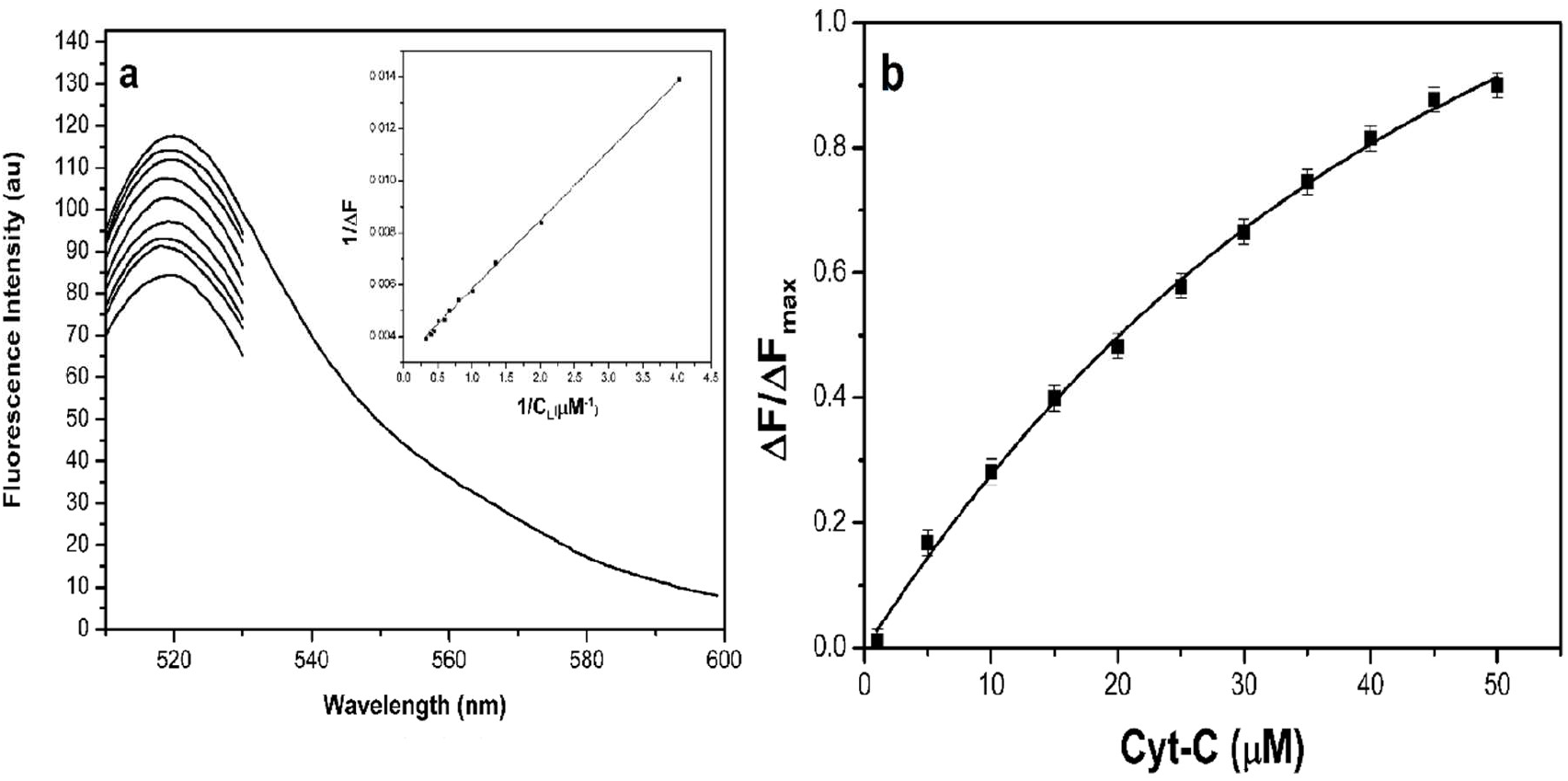
Panel ‘a’ shows the representative quenching curve of FITC labelled spectrin with cytochrome-c. F-spectrin in the range of 20-50 nM was excited at 495 nm and emission was monitored from 510-600 nm. Sequential addition of cytochrome-c from concentrated stock solution was done and resultant decrease in fluorescence intensity of F-spectrin was monitored at 520 nm. Inset shows the double reciprocal plot of the same from which ΔF_max_ is calculated. Panel ‘b’ shows the binding isotherm of the same from which K_d_ value is determined.

**Table 1:** The K_d_ values for the association of different heme proteins and spectrin is tabulated. Error bars represent S.D. of at least five independent experiments.

| Protein | K<sub>d</sub> (μM) |
| --- | --- |
| HbA | 27.6 ± 1.7 |
| HbA -α | 17.4 ± 1.5 |
| HbA - β | 25.3 ± 1.2 |
| HbE | 6.1 ± 1.6 |
| HbE -α | 16.2 ± 1.5 |
| HbE- β | 10.1 ± 1.4 |
| HbS | 21.6 ± 1.2 |
| HbS -α | 16.9 ± 1 |
| HbS- β | 20.2 ± 1.4 |
| Cytochrome-c | 0.89 ± 0.9 |

### Determination of liner range of enzyme activity and MM constants

The activity of spectrin was assayed in the concentration range of 10-40 nM and it was seen that spectrin does not possess detectable catalase or peroxidase activity. Hemoglobin used was checked for catalase activity due to the possible presence of contaminants, it was found that catalase activity was absent. The linear range and the K_m_ and V_max_ values of the enzyme activities of the heme proteins and catalytic activity of heme were determined and working concentrations were chosen accordingly. Representative plots of the linear range and Michaelis-Menten plots of HbA, HbE and HbS and their globin subunits are given in **Figure 2**. And those of catalase, cytochrome-c and heme are given in **Figure 3**. The MM plot of catalase could not be determined due to excessive bubbling under experimental conditions and that of heme could not be determined due to quick degradation in presence of peroxide. The values of the MM constants of the heme proteins and heme is tabulated in **Table 2.**

**Figure 2:**
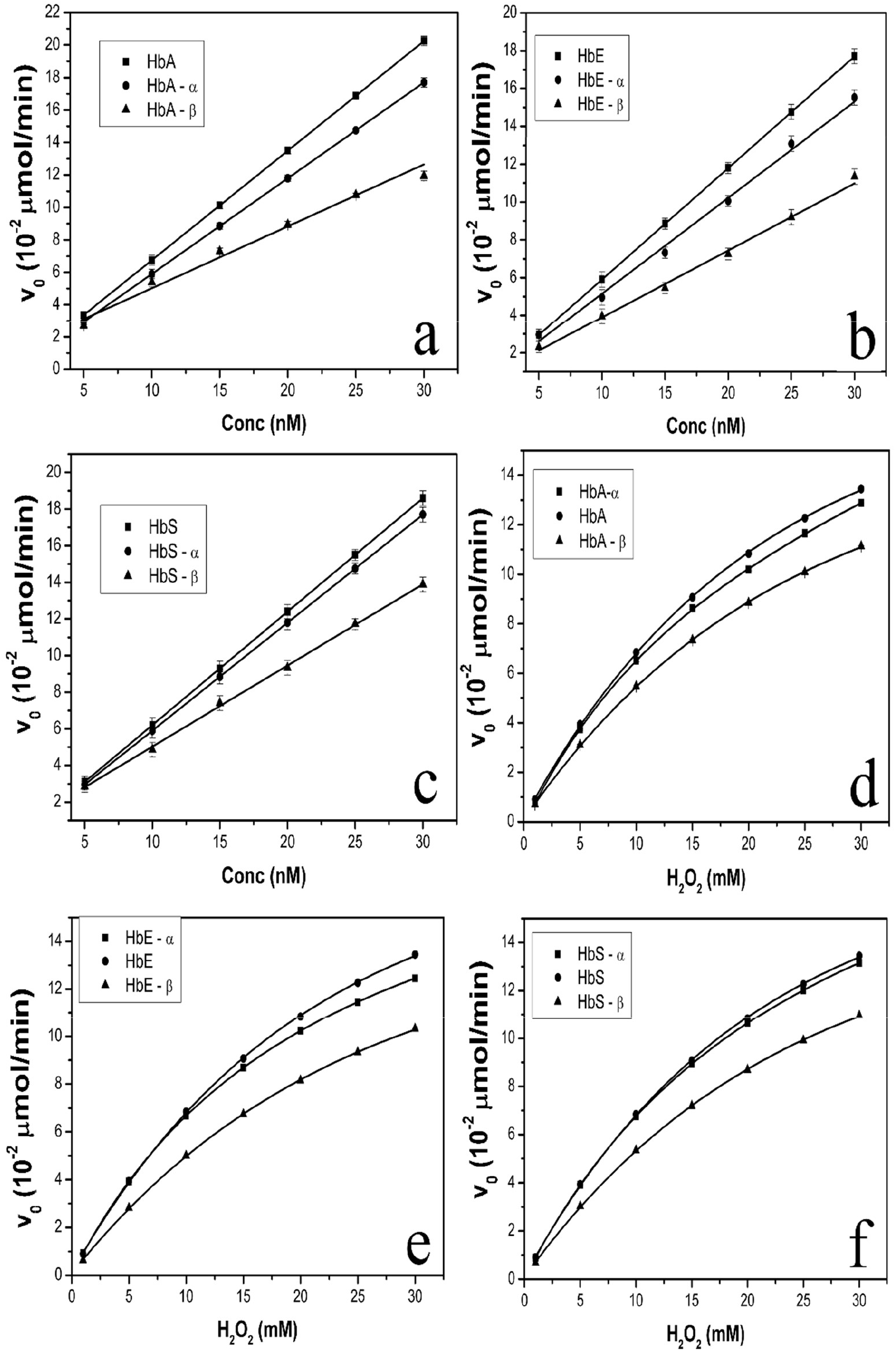
The linear range of hemoglobin variants and their isolated subunits as well as their Michaelis-Menten curves are shown. Peroxidase activity was measured in 50mM sodium-phosphate-citrate buffer pH 5.0 using 20 mM ABTS and 10 mM H_2_O_2_. The reaction was followed by monitoring the generation of coloured oxidation product of ABTS at 405 nm. The liner range of the peroxidase activity for all hemoglobin variants were determined from a plot of the initial velocity (v_0_) of the enzyme activity against the concentration of the enzyme. Panel ‘a’ shows HbA and its subunits, panel ‘b’ shows HbE and its subunits and panel ‘c’ shows HbS and its subunits. Using 20 mM ABTS and 10 nM of each of the respective hemoglobin variants and isolated globin subunits, the MM parameters were determined by plotting the initial velocity against peroxide concentration. Panel ‘d’ shows HbA and its subunits, panel ‘e’ shows HbE and its subunits and panel ‘f’ shows HbS and its subunits.

**Figure 3:**
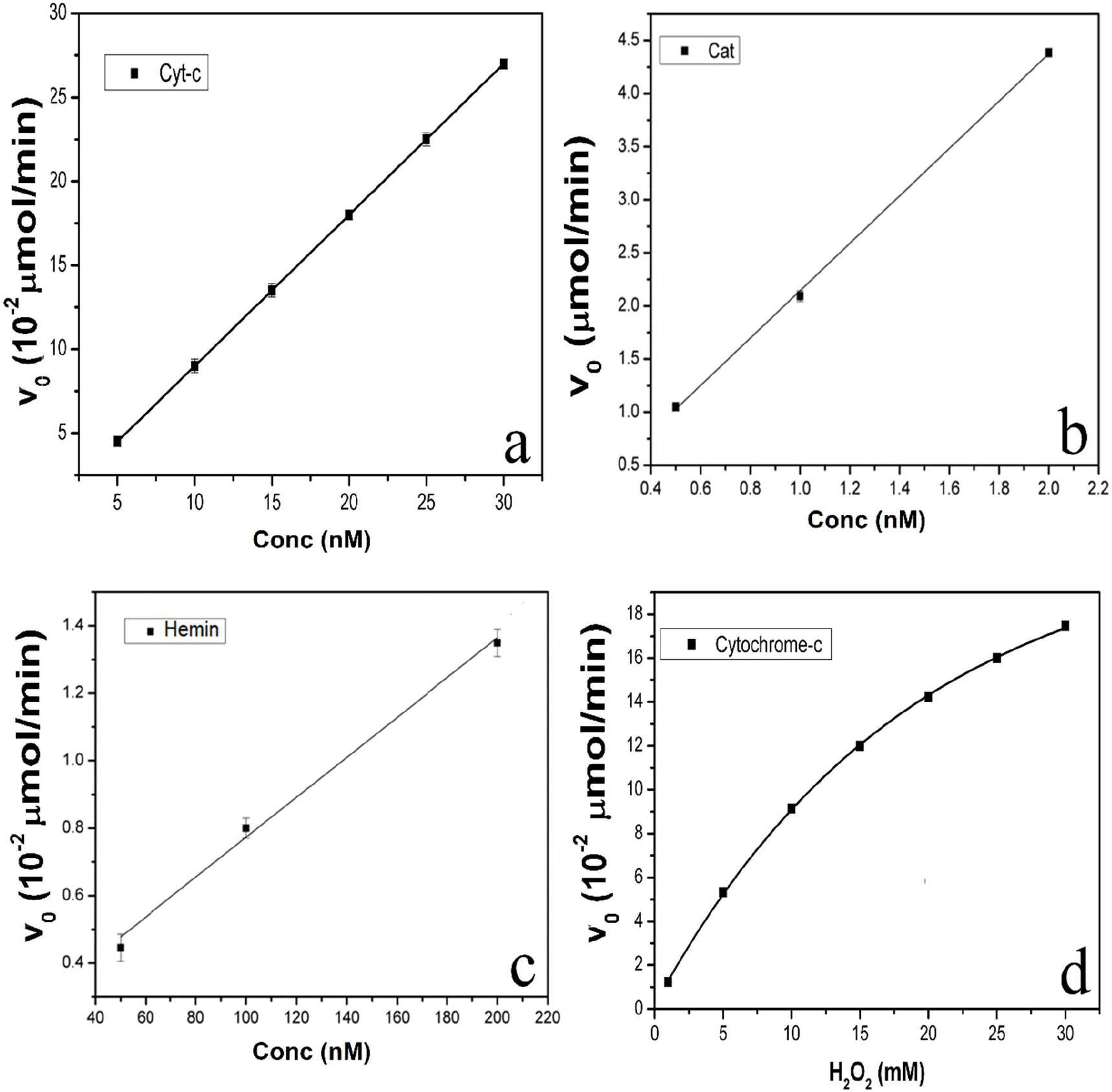
The linear range of cytochrome-c, catalase and heme (hemin) are shown. The liner range was determined from a plot of the initial velocity (v_0_) of the enzyme activity against the concentration of the enzyme. In case of catalase enzyme activity was assayed in 50mM Tris-HCl pH 8.0 buffer using 4 mM H_2_O_2_ and reaction was followed by monitoring the disappearance of peroxide at 240 nm. In case of cytochrome-c and heme (hemin), 50mM sodium-phosphate-citrate buffer pH 5.0 and 20 mM ABTS was used with 10 and 4 mM peroxide respectively. The reaction was followed by monitoring the generation of coloured oxidation product of ABTS at 405 nm. Panel ‘a’ shows linear range of cytochrome-c, panel ‘b’ shows that of catalase and panel ‘c’ shows that of heme (hemin). Using 20 mM ABTS and 10 nM cytochrome-c the MM plot was determined and is shown in panel ‘d’.

**Table 2:** The K_m_ and V_max_ values of the hemoglobin variants and their isolated subunits and cytochrome-c are listed. Values in parenthesis are those in presence of 20 nM spectrin. The error bars represent the S.D. of at least three independent experiments.

| Protein | K<sub>m</sub> (mM) | V<sub>max</sub> (10<sup>-2</sup> μmol/min) |
| --- | --- | --- |
| HbA | 30±1 (25±1) | 27±0.7 (33±0.5) |
| HbA -α | 28.2±1 (22±0.8) | 26±0.9 (32±0.6) |
| HbA - β | 32±0.9 (27±1) | 23±0.6 (28±0.4) |
| HbE | 23±0.9 (17±0.8) | 22±0.7 (29±0.8) |
| HbE -α | 28.4±1 (22±0.9) | 26±0.9 (32±0.5) |
| HbE- $\beta$ | 33±0.7 (29±1) | 23±0.5 (29±0.7) |
| HbS | 27±1 (22±0.8) | 25±0.7 (31±0.7) |
| HbS -α | 28.5±1 (23±1) | 27±0.9 (32±0.9) |
| HbS- $\beta$ | 34±0.8 (28±0.9) | 22±0.7 (29±0.4) |
| Cytochrome-c | 25±1 (20±1) | 32±0.5 (37±0.9) |

### Effect of spectrin on enzyme activity

Spectrin preparations were found to not possess any catalase or peroxidase activity of its own. Addition of BSA did not alter the enzyme activities of the heme proteins or catalytic activity of heme, showing that the modulation of enzyme activities seen on spectrin addition was specific to spectrin. It is seen that in general spectrin addition increased the enzymatic activity of all heme proteins under consideration and catalytic activity of heme in a dose dependent manner. The increase in enzyme activity in the presence of spectrin for hemoglobin variants and isolated globin chains is shown in **Figure 4**. The effect of spectrin on the activities of catalase, cytochrome-c, and heme is given in **Figure 5**.

**Figure 4:**
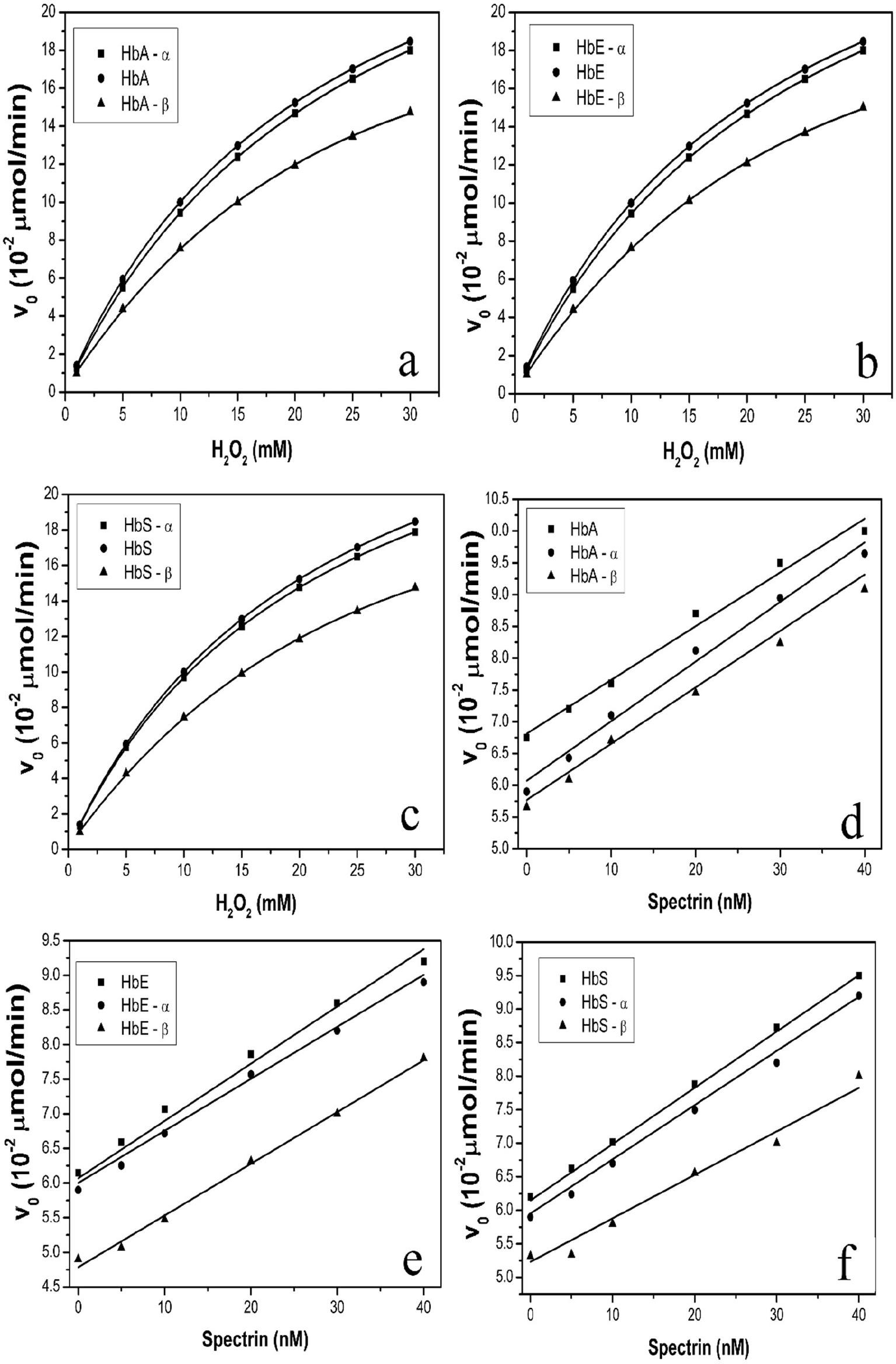
Using 10 nM of each hemoglobin variant and isolated globin subunit in presence of 20 nM of spectrin the MM curves of each were determined using 20 mM ABTS. Panel ‘a’ shows HbA and its subunits, panel ‘b’ shows HbE and its subunits and panel ‘c’ shows HbS and its subunits. The effect of increasing concentrations of spectrin on enzyme activity of hemoglobin was monitored using 10 nM of each hemoglobin variant and their isolated globin subunits using 10 nM of each protein, 10 mM peroxide and 20 mM ABTS. The initial velocity was plotted against spectrin concentration. Panel ‘d’ shows HbA and its subunits, panel ‘e’ shows HbE and its subunits and panel ‘f’ shows HbS and its subunits.

**Figure 5:**
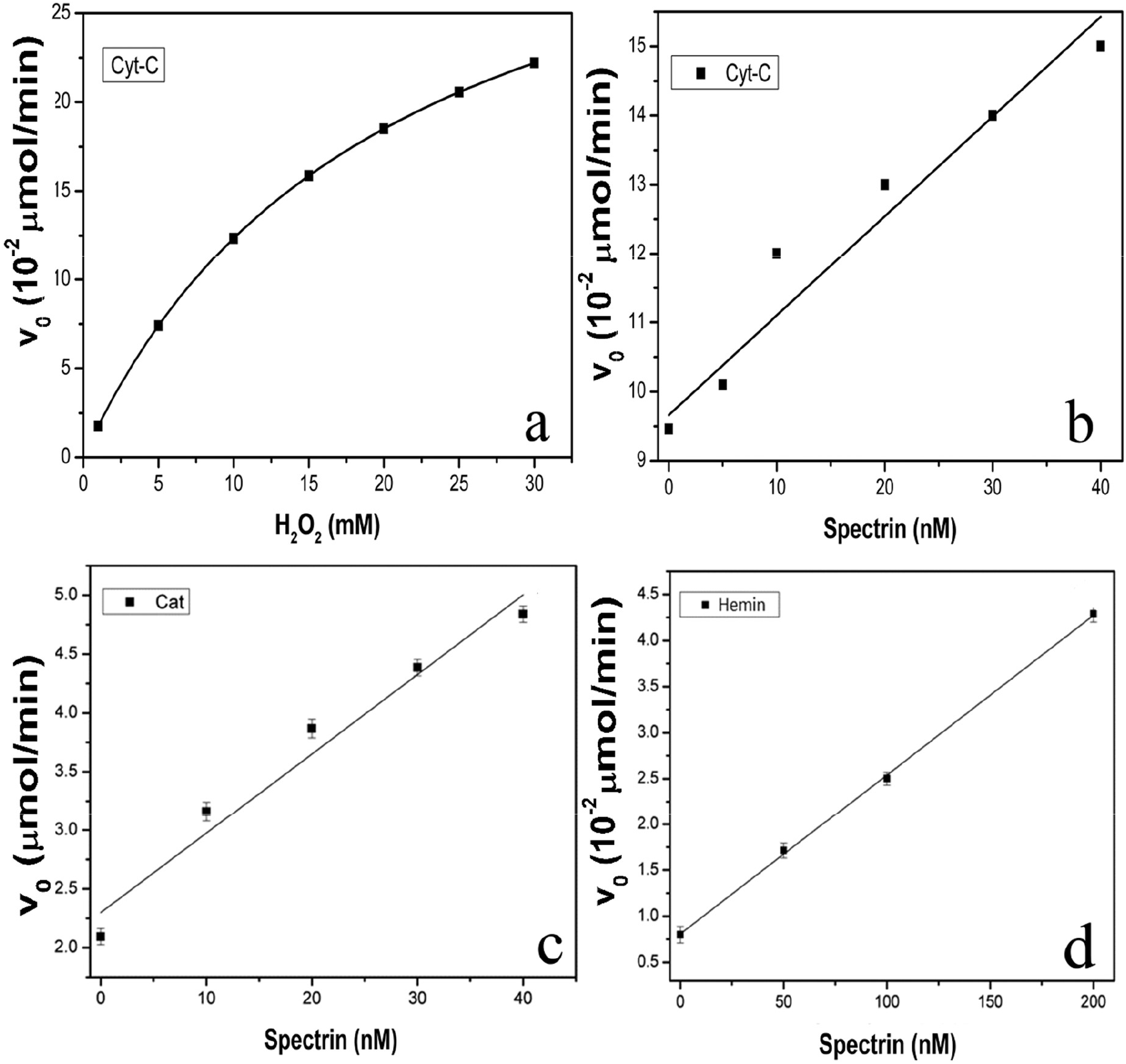
Panel ‘a’ shows the Mm curve of 10 nM cytochrome-c in presence of 20 nM spectrin and 20 mM ABTS. Panel ‘b’ shows the initial velocity of 10 nM cytochrome-c in presence of increasing amounts of spectrin and 10 mM peroxide and 20 mM ABTS. Panel ‘c’ shows the same for 1 nM catalase using 4 mM peroxide and 20 mM ABTS and panel ‘d’ for 100 nM heme (hemin) using 4 mM peroxide and 20 mM ABTS.

The values of the MM constants of the heme proteins and heme in presence and absence of 20 nM of spectrin is tabulated in **Table 2.**

It was seen that spectrin did not inhibit the degradation of heme proteins by peroxide, as measured from their residual activities. However, in case of heme addition of spectrin seemed to delay denaturation by hydrogen peroxide as evidenced by the greater residual activity with time in presence of spectrin versus its absence. It was also seen that under the experimental conditions spectrin formed cross-linked adducts with all the heme proteins except catalase. The extent of cross-linking was analysed according to the relative densities of band b, as in our previous work, and it was seen that HbE formed the highest amount of cross-linking followed by HbS and HbA and cytochrome-c [9]. Catalase did not form cross-linked products under experimental conditions.

We have previously shown the oxidative cross-linking of hemoglobin variants with spectrin [8, 9]. Here we show representative graphs of hydrogen peroxide mediated degradation of HbA and heme in presence of spectrin with respect to time, and the cross-linked aggregates of cytochrome-c, HbA, HbE and HbS in **Figure 6**.

**Figure 6:**
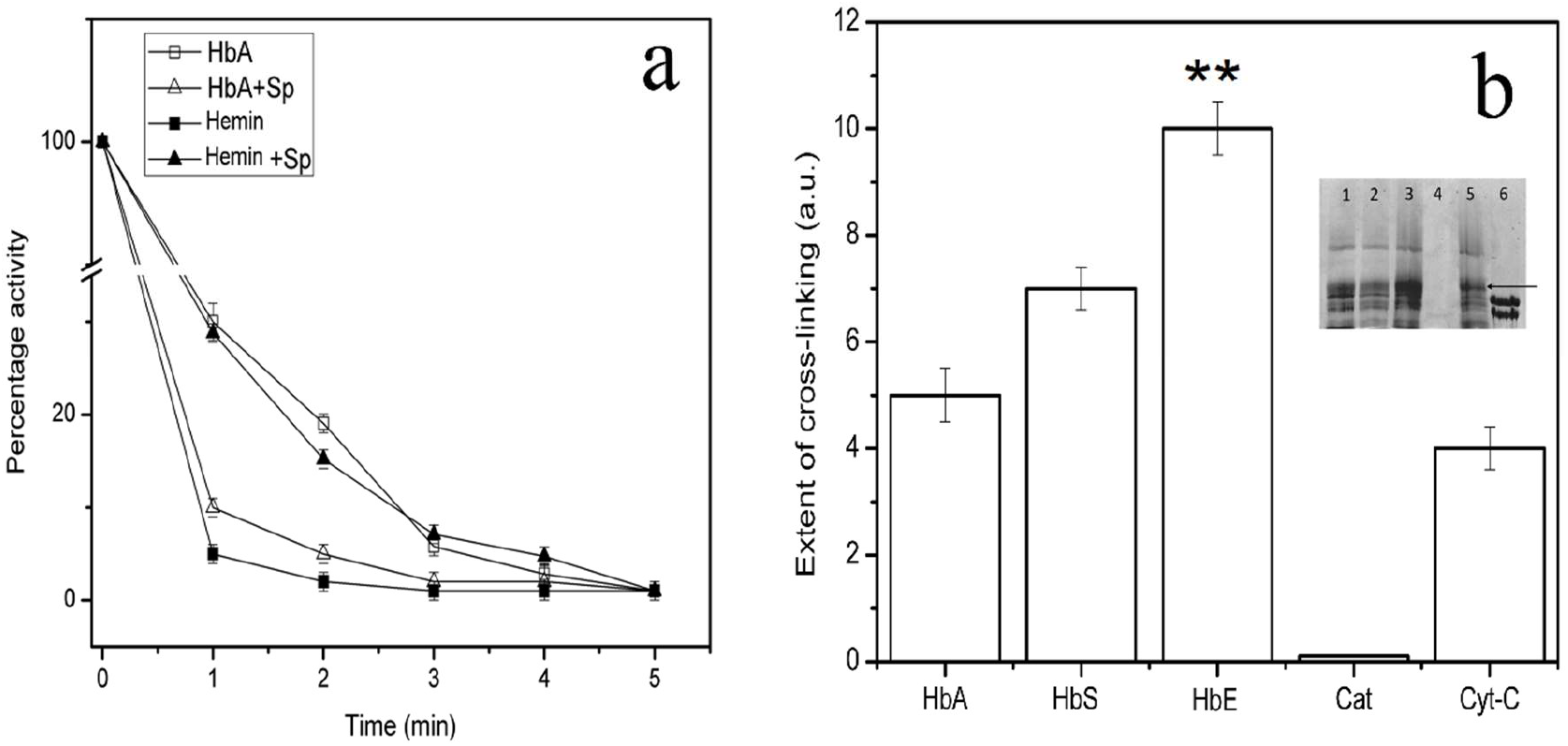
Panel ‘a’ shows the time dependent denaturation of 10 nM HbA and 100 nM heme (hemin) in 10 mM peroxide in presence and absence of 40 nM spectrin (Sp). HbA and heme in presence and absence of spectrin were incubated in peroxide for different times before ABTS was added and initial rate measured. Residual activity was expressed as percentage where native enzyme is considered 100%. Panel ‘b’ shows densitometric analysis of band b using ImageJ software. Error bars represent the S.D. of five independent experiments. Spectrin forms significantly greater amount of aggregates (as indicated by **) with HbE than other hemoglobin variants as tested by *t*-test with *p* values < 0.05. Inset shows 4% SDS gel of cross-linked aggregates of 1 μM each of heme proteins incubated with 1 μM spectrin in 4 mM peroxide at 37°C for 15 minutes. Lane 1 of inset shows HbS, lane 2 shows HbA, lane 3 shows HbE, lane 4 shows catalase, lane 5 shows cytochrome-c and lane 6 shows purified spectrin. The arrow represents the location of band b.

### Assay of lipid peroxidation

Extent of lipid peroxidation was assayed using the TBA method and quantifying the level of malondialdehyde (MDA) produced. It was seen that addition of heme proteins caused a significant increase in the production of MDA over blanks containing only SUVs and peroxide. Addition of ABTS with heme proteins reduced production of MDA and presence of spectrin along with it reduced it the most. Representative graphs for the TBA assay for HbA are given in **Figure 7.**

**Figure 7:**
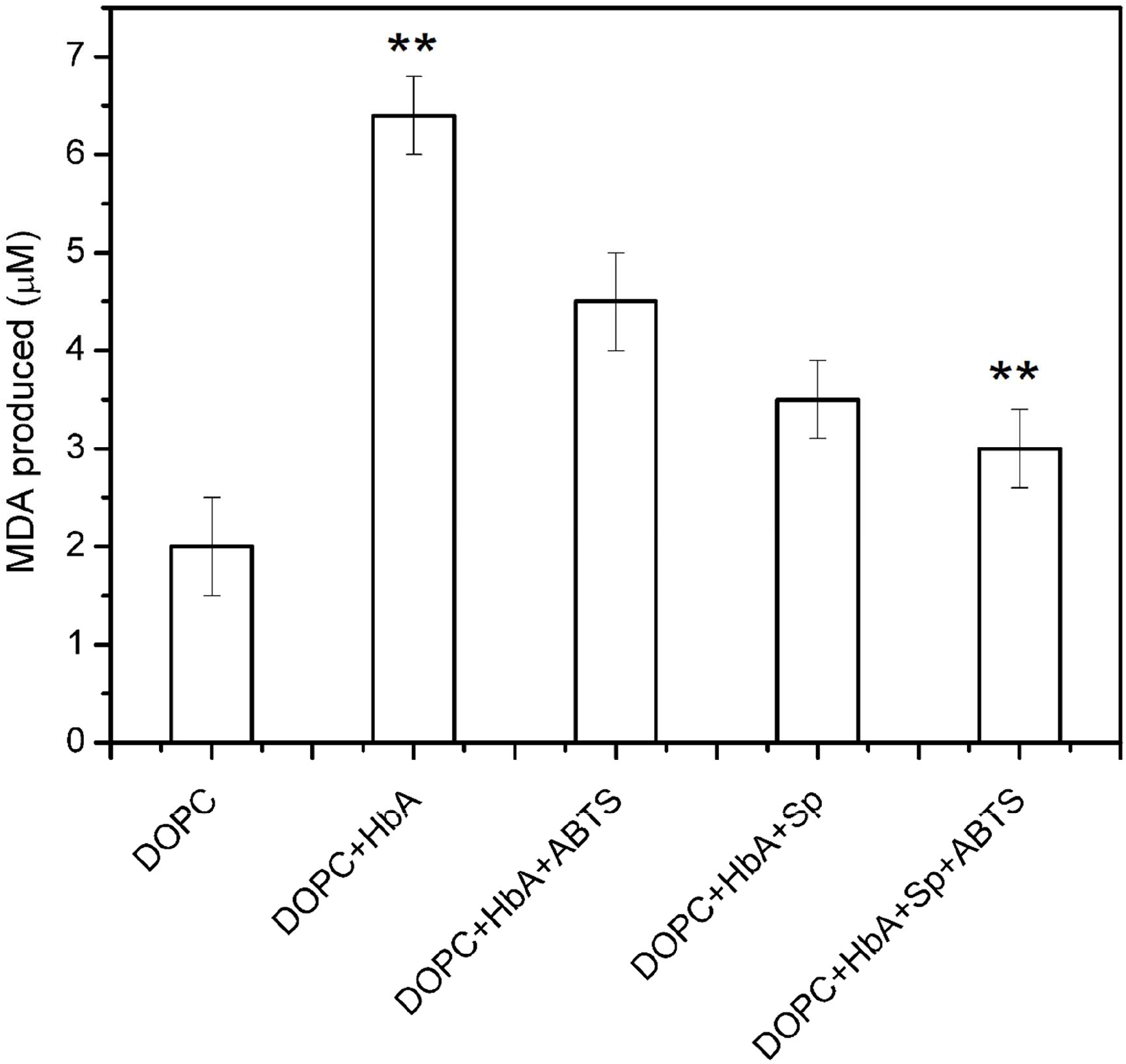
10 mM total phospholipid, of 100% DOPC SUVs were incubated for 15 minutes in 10 mM peroxide with or without 10 nM HbA in presence and absence of 40 nM spectrin (Sp) with or without additional 20 mM ABTS. From these mixtures, 0.2 μM total lipid was incubated with 2 ml of the TBA reagent in boiling water for 15 minutes and lipid peroxidation product malondialdehyde (MDA) was assayed at 535 nm. Error bars represent the S.D. of five independent experiments. Spectrin significantly reduced MDA formation as tested by *t*-test with *p* values < 0.05 (as indicated by **).

### Raman spectra of heme proteins and heme

The Raman spectra of hemoglobin variants was predominated by the spectra of their prosthetic heme group. Catalase did not show meaningful Raman spectra due to presence of auto-fluorescence. It was found that spectrin binding lead to a change in the spectra of the hemoglobin variants and cytochrome-c in a remarkably similar manner. It was seen that the peaks at positions 977 and 1004 cm^−1^ in all cases in absence of spectrin were overshadowed by a much more intense peak at 981 cm^−1^ in the presence of spectrin. In case of heme though there was no change seen in the position of the Raman peaks. However, the intensities of the peaks at 1065, 1371 and 1626 cm^−1^ were decreased with respect to peaks besides them in presence of spectrin than in their absence. Representative Raman spectras are shown in **Figure 8 & 9**.

**Figure 8:**
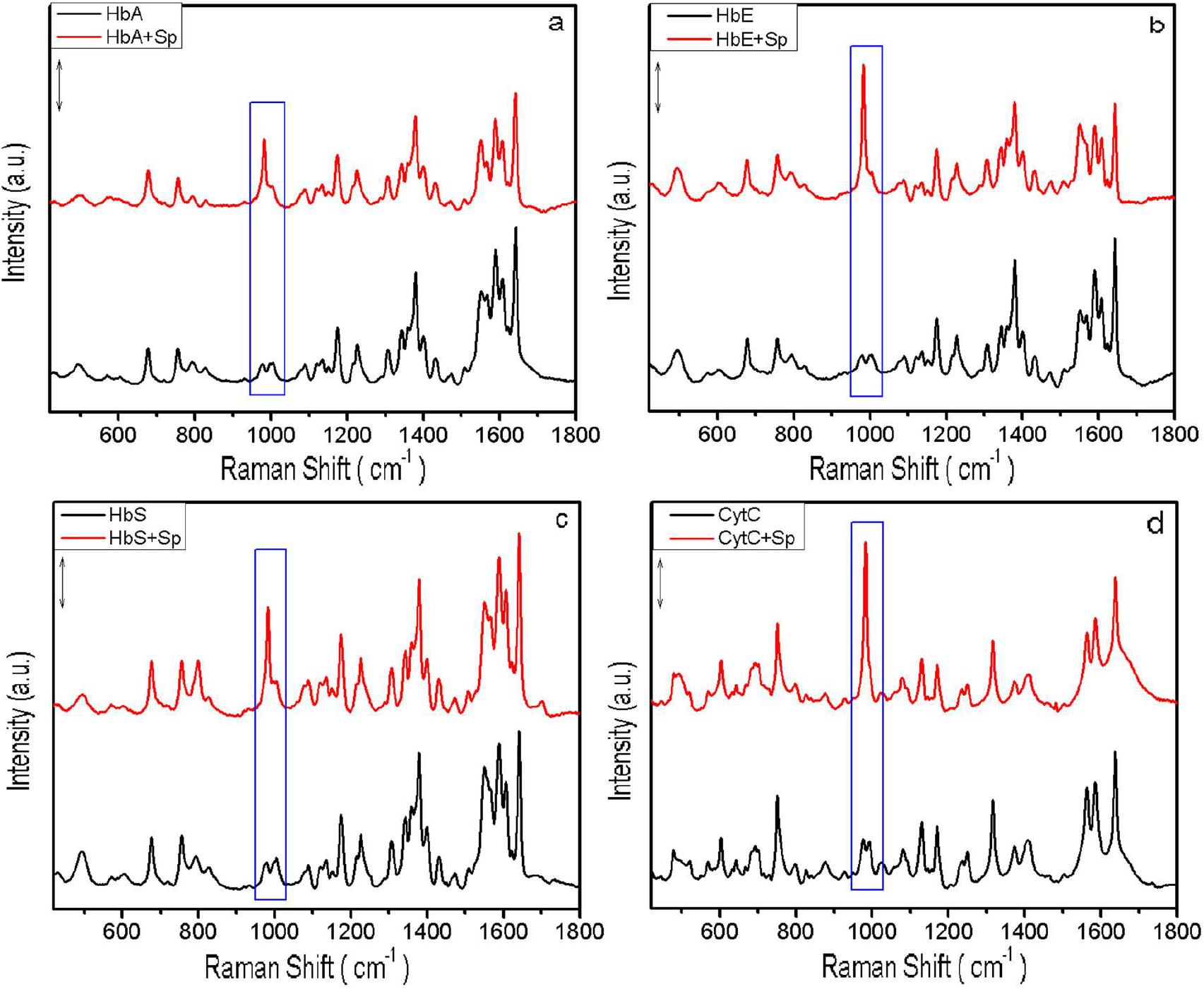
Panel ‘a’ shows the representative Raman spectra of 300μM HbA in the presence and absence of 3μM spectrin (Sp), panel ‘b’ shows the same for HbE, panel ‘c’ for HbS and panel ‘d’ for cytochrome c (CytC). The bars represent 2000 counts. The peaks where the major change is seen on spectrin binding are boxed.

**Figure 9:**
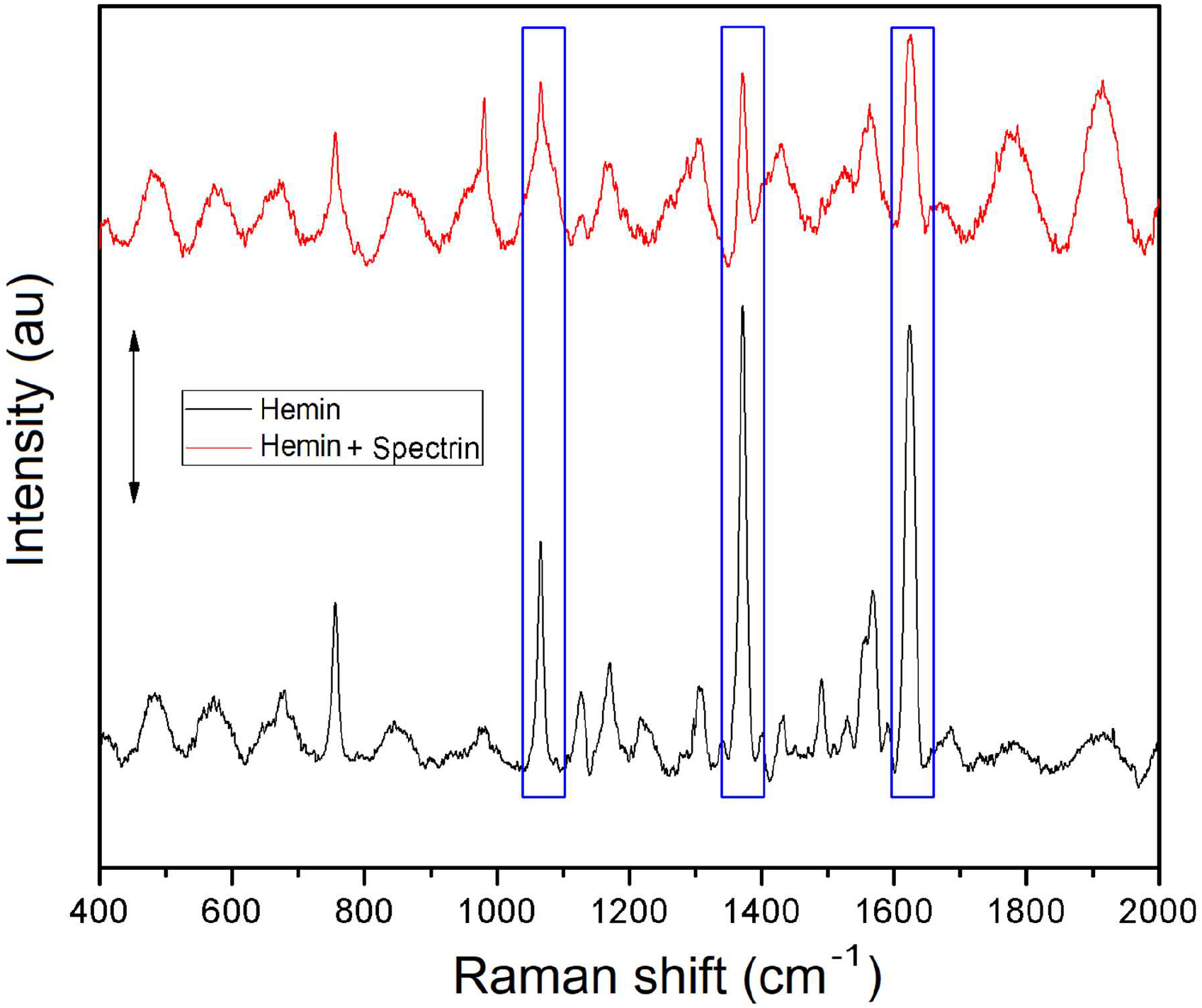
The representative Raman spectra of heme (hemin) in presence and absence of spectrin is shown. The bar represents 2000 counts. Peaks with the maximum change are boxed.

## Discussion

Our binding experiments with FITC-spectrin shows that spectrin interacts with all the heme proteins with comparable apparent equilibrium dissociation constants for the most part, establishing that spectrin is able to physically bind/interact with the target proteins.

Significantly it is seen that amongst the hemoglobins, spectrin interacts with the highest affinity to the most unstable/unstructured hemoglobin variant, HbE. This may have implications in disease states where the associated hemoglobin variant causes progression of the disease phenotype. HbE is the most unstable among the three variants investigated [18]. Previously we have shown the interaction between spectrin and HbA and HbE. This is the first report for the interaction of spectrin and HbS as well as a comparative study of spectrin interaction with three different hemoglobin variants and their isolated subunits.

It is interesting to note that spectrin interacts most strongly with cytochrome-c, a protein which is not found in the RBC but is mitochondria resident, moreover it shows the highest affinity for interaction with spectrin. This points to the fact that spectrin may interact with heme proteins in general; previous reports from our lab have also shown spectrin interaction with a non-RBC heme protein, HRP, which incidentally also showed spectrin involvement in HRP redox activity [10]. Here we show a novel role of spectrin in the redox activity of hemoglobins, catalase and cytochrome-c.

We see that the addition of spectrin increases the enzyme activities of all the heme proteins and the catalytic activity of heme. It is found that out of HbA, E and S the maximal increase is seen in case of HbE. This is in agreement with our observation that spectrin binds relatively unstable heme proteins with higher affinity than stable ones [2].

In the case of hemoglobin, the ability of spectrin to influence its redox activity becomes especially important considering that, oxidative stress inside an erythrocyte is a major challenge, especially so in the context of the RBC membrane where redox active hemes and hemoglobins give rise to many pathological membrane damages [42, 43]. The ability of spectrin to enhance peroxidase and catalase activity of heme proteins opens up a potentially novel way in which this oxidative challenge may be influenced *in situ*. In the presence of a suitable reducing substrate hemoglobin variants and globin subunits are capable of catalytically removing peroxide. The RBC cytosol has a high concentration of reducing substrates, namely glutathione and NADPH for the existent peroxidase machinery. Hemoglobin may also be able to utilize this substrate pool and reduce peroxide, thereby helping mitigate oxidative stress, and spectrin may help the process due to its ability to enhance peroxidase activity.

Similarly it has been shown that spectrin is a heme acceptor and can bind free heme with a dissociation constant of 0.57 ± 0.06 μM and mitigate its toxic effects [26]. Here we show that spectrin can enhance the peroxidase activity of free heme (hemin) as well, which may in turn help in fighting ROS challenge. In this regard it is worth pointing out that free heme when bound to polymers such as DNA or immobilized shows increased catalytic activity in general [44–46].

It is important to note that the promiscuous peroxidase activity of membrane bound hemoglobin and free heme is responsible for deleterious membrane lipid and protein oxidation. We have shown that spectrin may be able to mitigate this, as in cases where suitable peroxidase substrate is absent, heme proteins form covalently linked aggregates with spectrin and such aggregates are redox inactive. Moreover, the extent of cross-linking of spectrin with hemoglobin variants follows expected trends, HbA forms least cross-links, followed by HbS and HbE, which forms the most [9]. This inactivation of heme proteins on spectrin linkage must protect the membrane from further oxidative damage. In fact it is seen that on oxidative stress hemoglobins are found attached to spectrin *in vivo* [47].

It is known that haptoglobin, another chaperone binds cell free hemoglobin and the complex was hypothesised to decrease hemoglobin peroxidase activity, however, it was found that the complex retained its activity and consumed ascorbate as reductant [48]. This has similarities with our present work, with the difference that hemoglobin redox activity is lost on spectrin cross-linking.

Lipid peroxidation experiments lend support to this argument. It is seen that spectrin is able to decrease the amount of heme protein mediated lipid peroxidation in the absence of reducing substrates. Also due to its enhancing effect on enzyme activity, it is able to decrease lipid peroxidation to near baseline levels in presence of a reducing substrate.

Moreover, the ability of spectrin to enhance enzymatic activity extends to other heme proteins, namely catalase, which is a major player in the RBC defence against peroxide, thus spectrin can modulate the pre-existing cellular defence against ROS directly as well.

It must be kept in mind that the enhancement in enzyme activity that spectrin causes may come about either by protection against peroxide, or it by structural alteration upon binding that leads to an enhancement of catalase or peroxidase activity.

It is known that peroxide is a suicide substrate of peroxidases, thus agents that protect the enzyme from damage lead to a higher residual activity [47, 49]. However, it is demonstrated that incubation with peroxide in presence of spectrin affords no protection over that in its absence, thereby ruling out this possibility of spectrin protecting the enzyme from oxidative damage. In the case of heme though it is seen that residual catalytic activity is increased in the presence of spectrin, which may act by shielding the heme from peroxide.

Further, using Raman spectroscopy, we have shown that conformational alterations take place in hemoglobin variants, cytochrome-c and heme upon spectrin binding; and these in turn lead to increased enzyme activity. In case of hemoglobin variants and cytochrome-c it is seen that the peak at 977 cm^−1^ due the out of plane deformation mode of the C_a_-H bond and the peak at 1004 cm^−1^ due to phenylalanine get replaced by an out of plane deformation mode of the C_a_-H bond at 981 cm^−1^ [50–52]. This conformational alteration is seen across all the four proteins pointing to the fact that spectrin may interact with heme proteins in the same general way.

In case of heme though there is no appearance of new peaks, there is a decrease in the relative intensities of the peaks at 1065, 1371 and 1626 cm^−1^ which arise due to in plane deformation of C_b_-H_2_ bond, in plane symmetric stretching of pyrrole rings and in plane stretching of C_a_=C_b_ bond respectively [53]. It should be noted that this decrease may be due to the changed state (powder and solution) heme and heme - spectrin combination are investigated under as it may affect bond polarizability and concurrently peak intensity. However, as the relative intensities of these peaks with respect to adjacent peaks have decreased it points to probable conformational alteration.

It can be concluded that spectrin interacts with heme proteins and heme with apparent equilibrium dissociation constants in the low micromolar range indicating moderately strong binding. Spectrin binding enhances enzyme activity of heme proteins and heme which may help mitigate ROS challenge in the presence of reducing substrate. Further, spectrin heme protein cross-linking may abolish redox potential of these proteins which in turn may protect the cell membrane lipids and proteins from oxidative damage mediated by promiscuous activity in the absence of reducing substrate. It is shown that spectrin binding causes conformational changes in heme proteins and heme which are detectable by Raman spectroscopy and such changes are thought to give rise to enhancement in enzyme activity.

## Acknowledgement

The authors would like to acknowledge Dr. Debasis Banerjee of Hematology Division, Ramkrishna Misson Seva Prathisthan Hospital, Kolkata for kindly providing us with clinical samples.

## Abbreviations

ABTS: 2,2’-azino-bis(3-ethylbenzothiazoline-6-sulphonic acid
PMB: para-hydroxymercuribenzoic acid
DTT: di-thiothreitol
RBC: red blood cell
DEAE: diethylaminoethyl
CM: carboxymethyl
HbA: hemoglobin A
HbE: hemoglobin E
HbS: hemoglobin S
MDA: malondialdehyde
TCA: trichloro-acetic acid
TBA: thiobarbituric acid
MM: Michaelis-Menten

